# Daily locomotor rhythms and ecdysial stage-dependent behaviors in the tardigrade *Hypsibius exemplaris*

**DOI:** 10.64898/2026.09.23.753707

**Authors:** Baris Can Ulku, Lars Hering, Natalia Filimonova, Elfriede Friedmann, Georg Mayer

## Abstract

Tardigrades, key ecdysozoan organisms, have poorly characterized daily behavioral rhythms linked to their molting (ecdysial) cycle. We continuously tracked the locomotor activity of the model tardigrade *Hypsibius exemplaris* using infrared videography and deep learning-based automated tracking under three photic regimes: a light–dark cycle, constant darkness following entrainment, and constant darkness without prior entrainment. We identified a robust endogenous circadian-like rhythm in locomotor activity that persisted in constant darkness, independent of photic input. Although the light–dark cycle enhanced the rhythm amplitude and the overall animal activity, it was not required for rhythmicity. Individual phases varied widely, indicating a lack of population-level synchronization. Furthermore, we developed a behavioral classification system based on activity level, which enables categorizing distinct behaviors. Behavioral patterns were modulated by distinct ecdysial stages, with quiescence gating premolt and time-of-day dependent patterns in feeding and wandering. These findings provide evidence consistent with an endogenous, circadian-like timing system in tardigrades that integrates daily activity with molting cycles and establish *H. exemplaris* as a promising model for studying ecdysozoan chronobiology.

**Summary statement:** Tardigrade locomotor activity is shaped by an endogenous circadian-like timing system that persists without light input and is embedded within the ecdysial cycle. This work establishes *Hypsibius exemplaris* as a tractable model for studying how biological clocks interact with molting programs.

## Introduction

Organisms have evolved under recurring environmental cycles, driven primarily by daily fluctuations in light and temperature. These periodic external cues, or zeitgebers, synchronize endogenous biological clocks that regulate behavior, physiology, and metabolism (Aschoff, 1960). The most prominent of these timing mechanisms is the circadian clock—a self-sustained oscillator that generates approximately 24-hour rhythms and is entrained by shifting environmental signals. Circadian networks are remarkably ubiquitous, spanning bacteria (Wollmuth and Angert, 2023), fungi (Dunlap et al., 2007), plants (Dodd et al., 2005), and animals (Wood et al., 2020). Comparative studies have revealed both deeply conserved molecular principles and extraordinary structural diversity, highlighting how biological timing has adapted to diverse evolutionary pressures (Bell-Pedersen et al., 2005). To understand the evolutionary history of these systems, it is critical to investigate organisms occupying key phylogenetic nodes. Tardigrades represent one such lineage, constituting the sister group to Arthropoda and Onychophora (Giacomelli et al., 2025). Yet, despite their evolutionary importance, circadian rhythmicity remains unexplored in this clade at both the behavioral and molecular levels.

Tardigrades (water bears) are microscopic metazoans belonging to the Ecdysozoa, a superclade unified by the physiological process of molting (Aguinaldo et al., 1997; Fleming and Arakawa, 2021). Despite their anatomical and genomic simplicity (Gross et al., 2019), recent behavioral studies have revealed that tardigrades possess a surprisingly sophisticated behavioral repertoire. This includes targeted survival responses to environmental extremes (Jönsson, 2019; Hagelbäck and Jönsson, 2023), active mate-searching strategies (Chartrain et al., 2023), complex, coordinated locomotion (Nirody et al., 2021; Anderson et al., 2024), intricate prey–predator interactions (Meyer et al., 2020), and associative learning (Zhou et al., 2019). By contrast, their responses to light remain poorly understood, and the available evidence is historically contradictory: early work suggested age-dependent photokinesis in *Macrobiotus hufelandi* (Beasley, 2001), whereas a subsequent study reported no clear light– dark effects on the activity of *Milnesium tardigradum* (Shcherbakov et al., 2010). Crucially, no study has systematically investigated how behavioral states are organized across the broader ecdysial cycle, leaving the relationships between light input, molting state, and daily timing unresolved.

We selected *Hypsibius exemplaris* as a model system to address these gaps because it reproduces parthenogenetically, enabling uniform clonal lines that minimize genetic variability, and has a well-annotated genome (Yoshida et al., 2017; Goldstein, 2018). Adult individuals are exceptionally small (∼200 μm), and their molting cycle is tightly coupled to oviposition throughout most of their lifespan (Nelson, 2002; Goldstein, 2018; Stone and Vasanthan, 2020). *H. exemplaris* is also an ideal candidate for photic timing research, as it possesses multiple opsin proteins and distinct photosensory cell architectures (Hering and Mayer, 2014; Gross et al., 2019; Fleming et al., 2021). Notably, non-visual opsins—molecules that frequently drive circadian photoreception in other animals (Buhr and Van Gelder, 2026)— are expressed across widespread peripheral tissues in this species, including the brain, ventral trunk ganglia, midgut, and storage cells (Dutta et al., 2026 preprint). Together, these features provide a strong physiological basis for testing whether daily locomotor activity is modulated by interactions between endogenous timing signals and light input.

Currently, rhythmic gene expression and canonical molecular clock components remain unverified in tardigrades. Although three pigment-dispersing factor genes (*He-pdf-1*, *-2*, and *-3*) have been identified in *H. exemplaris* (Mayer et al., 2015; Dutta et al., 2025), their precise roles as circadian coupling factors have yet to be functionally demonstrated. Consequently, characterizing the fundamental behavioral baseline of daily rhythmicity is a critical prerequisite for future molecular inquiries into candidate clock genes. Here, we establish a robust framework for long-term behavioral tracking in *H. exemplaris* to quantify how specific behavioral states fluctuate across ecdysial transitions and varying photic regimes. While further investigations into phase resetting and temperature compensation will be required to confirm a canonical circadian clock (Vitaterna et al., 2001), this work provides evidence consistent with endogenous, circadian-like modulation of activity in tardigrades and establishes a new entry point into ecdysozoan chronobiology.

## MATERIALS AND METHODS

### Tardigrade husbandry and culture conditions

Specimens of *Hypsibius exemplaris* Gąsiorek, Stec, Morek & Michalczyk, 2018 (Eutardigrada, Hypsibiidae; strain Sciento Z151; McNuff, 2018) were maintained as a clonal line derived from a single egg. Cultures were housed in polystyrene Petri dishes (100 mm × 20 mm; Sarstedt, Nümbrecht, Germany) containing Volvic mineral water (Danone Waters Deutschland GmbH, Frankfurt am Main, Germany) supplemented with *Chlorococcum* sp. algae as the sole food source. Animals were reared at a constant room temperature of 20 ± 2 °C under either a standard 12 h:12 h light–dark cycle (LD; lights on at 08:00, off at 20:00) or constant darkness (DD), depending on the experimental cohort. Culture media and food supply were refreshed biweekly to maintain stable environmental conditions and ensure age-synchronized populations. To minimize behavioral variability associated with molt-dependent state, single individuals actively undergoing molting were isolated for behavioral trials.

### Custom arena architecture

To facilitate long-term, high-resolution tracking without substrate degradation, we developed a custom agarose arena system. A 1–2 mm layer of molten 1% pulsed-field agarose (Carl Roth GmbH + Co. KG, Karlsruhe, Germany)—selected for its structural stability and resistance to dissolution during extended aquatic recordings—was poured into a 100 mm × 20 mm Petri dish at 60 °C. The agarose was allowed to partially solidify for ∼1 min. An acrylic glass tube (polymethyl methacrylate; 5 mm inner diameter × 8 mm height; Asia Pacific Elite Ltd, Kwai Fong, Hong Kong) was then positioned vertically onto the semi-solidified agarose base (Fig. 1A). Following complete solidification (∼30 min), the dish was rotated prior to imaging to minimize shadows over the edge of the arena. Between experimental replicates, acrylic tubes were thoroughly cleaned with 70% ethanol, rinsed with distilled water, and dried under gentle air pressure. Tube reuse was assessed to confirm that repeated use did not affect tracking efficiency or baseline animal behavior.

**Fig. 1.**
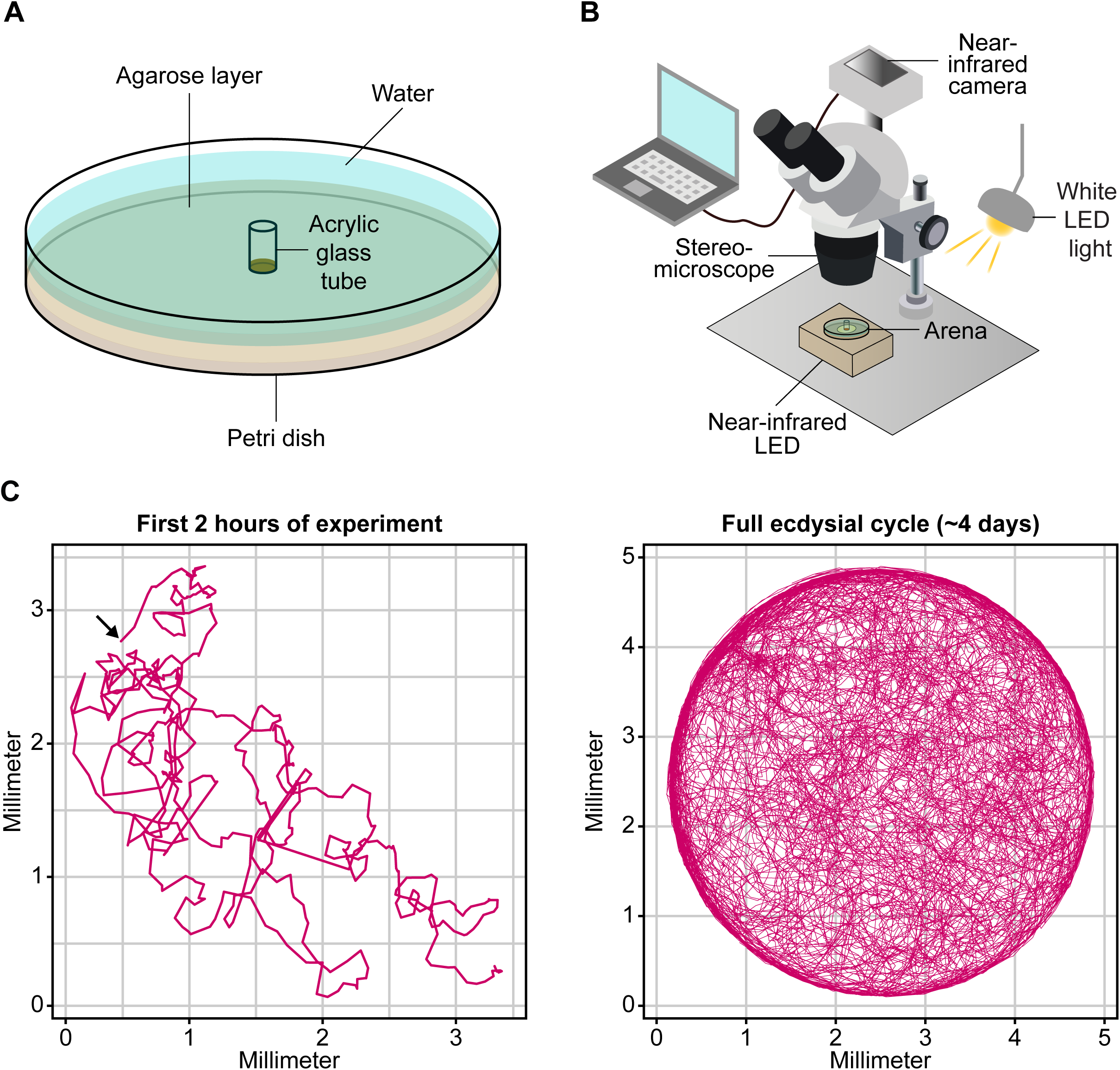
Experimental setup for long-term locomotor tracking of *Hypsibius exemplaris*. (A) Schematic of the custom behavioral arena. A 1–2 mm layer of 1% pulsed-field agarose lines the bottom of a 100 mm × 20 mm Petri dish. A cylindrical PMMA confinement chamber (5 mm diameter × 8 mm height) is positioned on the substrate. The arena is filled with Volvic water. (B) Imaging configuration. Top-down illumination is provided by a white LED (30 lux), and near-infrared (NIR) illumination (Urceri MT-912) is mounted beneath the arena. Imaging is captured through an Olympus SZ61 stereomicroscope (45× magnification) equipped with an NIR-sensitive camera. (C) Representative locomotor activity trace from a single animal, binned at 12-s intervals. The left panel displays the first 2 h of recording (start position indicated by arrow); the right panel shows the complete recording period.

### Behavioral recording setup

Locomotor behavior was recorded using a custom stereomicroscope-based optical imaging pipeline (Olympus SZ61, Olympus Corporation, Tokyo, Japan; Fig. 1B, C; Movies S1–S4). Illumination for the light phases of the trials was provided by a top-mounted white LED with an intensity of 30 lux, as measured at the arena level using a luxmeter (Urceri MT-912, Shenzhen Flus Technology Co. Ltd, Shenzhen, China). To capture behavior continuously during dark phases without disrupting light-sensitive pathways, near-infrared (NIR) imaging was conducted using a monochrome camera (Manta G-419B NIR, Allied Vision Technologies GmbH, Stadtroda, Germany) paired with an independent NIR illumination source (IR7373C; Everlight Electronics Co. Ltd, Taipei, Taiwan) mounted beneath the arena floor (Fig. 1B). Automated image acquisition was controlled via µManager (Edelstein et al., 2010), integrated with ImageJ (Schneider et al., 2012) for video compilation. All recording trials were conducted at a temperature of 20 ± 2 °C.

### Light regimes and experimental design

Animals were assigned to one of three distinct photic regimes:

- **LD (Light–Dark Control):** Animals were both reared and recorded under a cycling 12 h:12 h light–dark schedule.
- **DD (Entrained Constant Darkness):** Animals were reared under a 12 h:12 h LD cycle to achieve environmental entrainment, then transferred to and recorded under constant darkness.
- **Naïve-DD (Naïve Constant Darkness):** Animals were maintained in absolute darkness from the egg stage onward and recorded exclusively under constant darkness.

Prior to testing, animals underwent a standardized feeding phase in a fresh glass dish containing Volvic water and *Chlorococcum* sp. algae (2 days for LD and naïve-DD; 3 days for DD cohorts). Following feeding, individual tardigrades were introduced via micropipette into the acrylic arena tube, which was filled with Volvic water and lightly coated with algae across the agarose surface. The stereomicroscope magnification was fixed at 45×, and image capture commenced immediately. For LD trials, the daytime arena-level light intensity was maintained at 30 lux. For both DD and naïve-DD conditions, all visible light sources were physically shielded, and the recording environment was maintained in constant darkness. Automated time-lapse images were captured at 12-s intervals (5 frames min^−1^) continuously for 4–7 days per animal.

### Deep learning video tracking

Locomotor trajectories were extracted using the multi-animal framework of DeepLabCut (DLC-ma, v2.3.10; Mathis et al., 2018). A custom training dataset comprising 17 representative videos (containing 3–12 animals each) was generated by manually labeling five anatomical landmarks: *head*, *trunk*, *back*, *left leg*, and *right leg*. To ensure robust tracking independent of minor body contortions, only the coordinates of the central trunk landmark were used for downstream kinematic analyses. The model was trained for 200,000 iterations using a ResNet-50 architecture with default parameters. Tracking accuracy was visually validated using the “Make Labeled Video” subroutine (Movie S1). Spatial coordinate refinement and manual corrections of tracklets were performed during the initial hours of recording using the “Refine Tracklets” tool, specifically to correct for low-contrast tracking failures characteristic of early postmolt animals.

### Kinematic data post-processing

Raw pixel coordinates were processed in R (v4.5.2, 2025; https://www.R-project.org/) using custom scripts derived from the DLCAnalyzer package (Sturman et al., 2020). Outlier frames demonstrating a landmark likelihood threshold < 0.1 or an instantaneous spatial displacement > 100 µm were discarded and replaced using linear interpolation. To isolate true displacement from video noise, an animal was classified as actively “moving” only if its instantaneous velocity exceeded 5.4 µm·s⁻¹ (the minimum sustained baseline velocity determined in pilot characterizations) for a continuous duration of at least 2 min (10 consecutive frames). Movements falling below these combined velocity and duration thresholds were filtered out as static tracking noise.

Locomotor tracking data were integrated into cumulative 10-min bins. Bins associated with prolonged stationary resting or localized feeding on algal clusters were visually audited and zero-corrected if displacement was entirely attributable to localized vertical crawling within the algal matrix. Activity metrics were normalized to the mean distance covered across all non-zero active bins within the intermolt stage. To eliminate nonstationary trends across the ecdysial cycle, each 10-min binned time-series dataset was detrended using the *zoo* package (Zeileis and Grothendieck, 2005) in R by subtracting a centered rolling median baseline calculated over a 30-h moving window. This mathematical filtering preserved ∼24-h rhythmic periodicities while isolating detrended residuals for subsequent rhythmicity analysis and circadian time (CT) waveform construction.

### Ecdysial staging and behavioral annotation

The ecdysial cycle was aligned with the 10-min activity bins. Ecdysial stages were defined according to explicit visual, morphological, and behavioral criteria:

- **Intermolt (INT):** Begins immediately after complete emergence from the old cuticle and is characterized by alternating bouts of exploratory wandering and active feeding.
- **Premolt (PRM):** Begins with the onset of behavioral quiescence and is marked by the cessation of both forward locomotion and feeding activity.
- **Molt (MLT):** Begins when the animal visibly detaches its body wall from the old outer cuticle and ends upon complete emergence from the old cuticle. During this stage, observed behaviors are restricted primarily to cuticle shedding and oviposition.

Specific behavioral states were assigned to each 10-min interval. Because zero locomotor activity could indicate feeding, resting, or cuticle shedding/oviposition (the “three zeros” problem), we distinguished these behavioral states using the following kinematic criteria:

- **Wandering:** Cumulative distance > 3.5 mm sustained for ≥ 20 min during the INT stage (Movie S1).
- **Feeding:** Cumulative distance < 3.5 mm during the INT stage, reflecting slow, localized locomotion on the arena floor or browsing within algal clusters (Movie S2).
- **Resting:** Cumulative distance = 0 mm during the PRM stage, indicating complete behavioral quiescence interrupted only by occasional internal muscle contractions (Movie S3).
- **Cuticle shedding/oviposition:** Assigned to all time bins corresponding to the MLT stage (Movie S4).

### Daily activity profile visualization

To visualize population-level time-of-day dynamics, individual activity and behavioral profiles were aggregated into hourly zeitgeber time (ZT) intervals, with ZT0 defined as the onset of the light phase. For LD cohorts, ZT was calculated directly from the physical light cycle. For DD, a projected ZT timeline was extrapolated from the animals’ preceding light–dark schedule. For locomotor activity profiles, median hourly activity profiles were calculated for each individual, and the collective median across all individuals was plotted to yield population-level profiles, thereby ensuring equal weighting across distinct experimental subjects. For behavior timing profiles, total time spent in each annotated behavior was summed hourly for each individual. Population-level behavioral profiles were then computed as the median of these per-individual cumulative times at each ZT. Therefore, y-axis scales were allowed to vary across behaviors.

### Statistical framework

#### Photic preference and time-of-day dependence

Populations were evaluated for light–dark phase preference using paired Wilcoxon signed-rank tests comparing median intermolt activity between the subjective light (ZT0–ZT11) and dark (ZT12–ZT23) phases. One-sided tests were applied to isolate directional trends. To statistically evaluate population-level time-of-day modulation, cyclic generalized additive mixed models (GAMMs) were fitted using the *mgcv* package (Wood, 2004) in R. The response variable was log-transformed activity: *log*(1 + *y*), where y is the normalized activity in each 10-min bin. ZT was incorporated as a cyclic cubic regression spline with a fixed 24-h periodicity (bs = “cc”, knots at 0 and 24 h), and individual identity was included as a random intercept (bs = “re”). To determine whether the temporal distribution of discrete behavioral states (wandering, feeding, resting, cuticle shedding/ovipositon) varied significantly across the day, repeated-measures ANOVA (RM-ANOVA) and non-parametric permutation tests (5,000 permutations) were performed to test for deviations from temporal uniformity.

#### Rhythmicity detection and phase analysis

Circadian rhythmicity was evaluated using hourly binned, detrended activity-tracking data restricted to the initial 60 h of the highly active INT stage. A fixed-period (24-h) cosinor model was fitted separately to each individual trajectory. The model was defined as: *y(t) = M + b_1_ cos (2πt/24) + b_2_ sin (2πt/24) + ε(t)*, where M represents the mesor (rhythm-adjusted mean), b_1_ and b_2_ are the cosine and sine regression coefficients, respectively, and ε(t) denotes the residual error. Cosinor amplitude was derived from the fitted coefficients as: 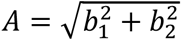. Goodness-of-fit was assessed using the coefficient of determination (*R²*), which quantifies the proportion of variance in the observed data explained by the fitted model. Rhythmicity was additionally assessed using the zero-amplitude F-test of the fitted 24-h cosinor model against a flat null model, testing the null hypothesis that *b_1_* = *b_2_* = 0. Because adjacent time-series observations are autocorrelated, *R²* was interpreted as a descriptive effect-size measure rather than a standalone formal test statistic. An *R²* threshold of 0.08 was additionally applied to identify trajectories with detectable circadian structure, and sufficiently stable acrophase estimates for group comparisons. Differences in absolute activity levels, amplitude (*A*), and rhythm strength (*R²*) across lighting groups were evaluated using Kruskal-Wallis tests. Population-level clustering of acrophases was assessed using the Rayleigh tests for non-uniformity of circular distributions, implemented in the R package *circular* (https://CRAN.R-project.org/package=circular).

#### Circadian time (CT) waveform realignment

To separate the common circadian waveform from inter-individual phase differences, activity trajectories of rhythmic animals were phase-aligned to an internal circadian time (CT) scale. For each individual, the cosinor-estimated acrophase was defined as CT0. Detrended 10-min intermolt activity residuals were assigned to their corresponding CT modulo (0–24 h), and individual waveforms were constructed using 30-min CT bins. Group-level median waveforms were subsequently derived across all rhythmic individuals within each experimental condition. To assess the long-term persistence of these endogenous patterns, the realignment pipeline was extended to generate continuous 48-h CT waveforms (Cycle 1: 0–24 h; Cycle 2: 24–48 h).

#### Resampling bootstrap waveform metrics

To statistically compare the underlying circadian waveform shape and peak-to-trough amplitudes across environmental groups, an animal-level bootstrap resampling procedure was performed (B = 2,000 iterations). In each bootstrap replicate, individual animals were resampled with replacement, and the group-level median CT waveforms were recomputed. Pearson correlation coefficients (*r*) between condition pairs were calculated to quantify structural waveform similarity. Waveform modulation amplitude, defined as maximum peak-to-trough distance of the median CT waveform, was calculated for each bootstrap replicate. Point estimates and robust 95% bootstrap confidence intervals (CIs; derived from the 2.5^th^–97.5^th^ percentiles) were reported for all cross-condition comparisons.

#### Software, panel design, and AI disclosures

All data visualizations and statistical curves were plotted using the *ggplot2* package (Wickham, 2009) in R. Behavioral curves were smoothed using locally estimated scatterplot smoothing (LOESS; smoothing parameter span = 0.03) implemented with the *ggplot2* package. Final multipanel figures were assembled and formatted in Adobe Illustrator CS5.1 (Adobe, San Jose, CA, USA).

AI-assisted language tools (ChatGPT, OpenAI) were used solely to refine grammar, syntax, and style. The authors independently reviewed and verified all technical content, data interpretations, and statistical analyses and assume full responsibility for the accuracy, transparency, and integrity of the presented work.

## RESULTS

### Ecdysial cycle progression dictates behavioral states

To characterize the baseline temporal organization of behavior, we continuously tracked individual *Hypsibius exemplaris* for 4–7 days across three distinct photic regimes: (1) a 12 h:12 h light–dark cycle (LD); (2) constant darkness following LD entrainment (DD); and (3) constant darkness without prior environmental entrainment (naïve-DD). High-resolution locomotor trajectories sampled at 12-s intervals were aggregated into 10-min and 1-h bins. Supervised behavioral classification identified four primary, mutually exclusive behavioral states: *wandering*, *feeding*, *resting*, and *cuticle shedding/oviposition* (Table 1; Movies S1–S4). The composition and temporal progression of these behavioral states follow a highly stereotyped, recurring sequence across ecdysial cycles (Fig. 2A).

**Fig. 2.**
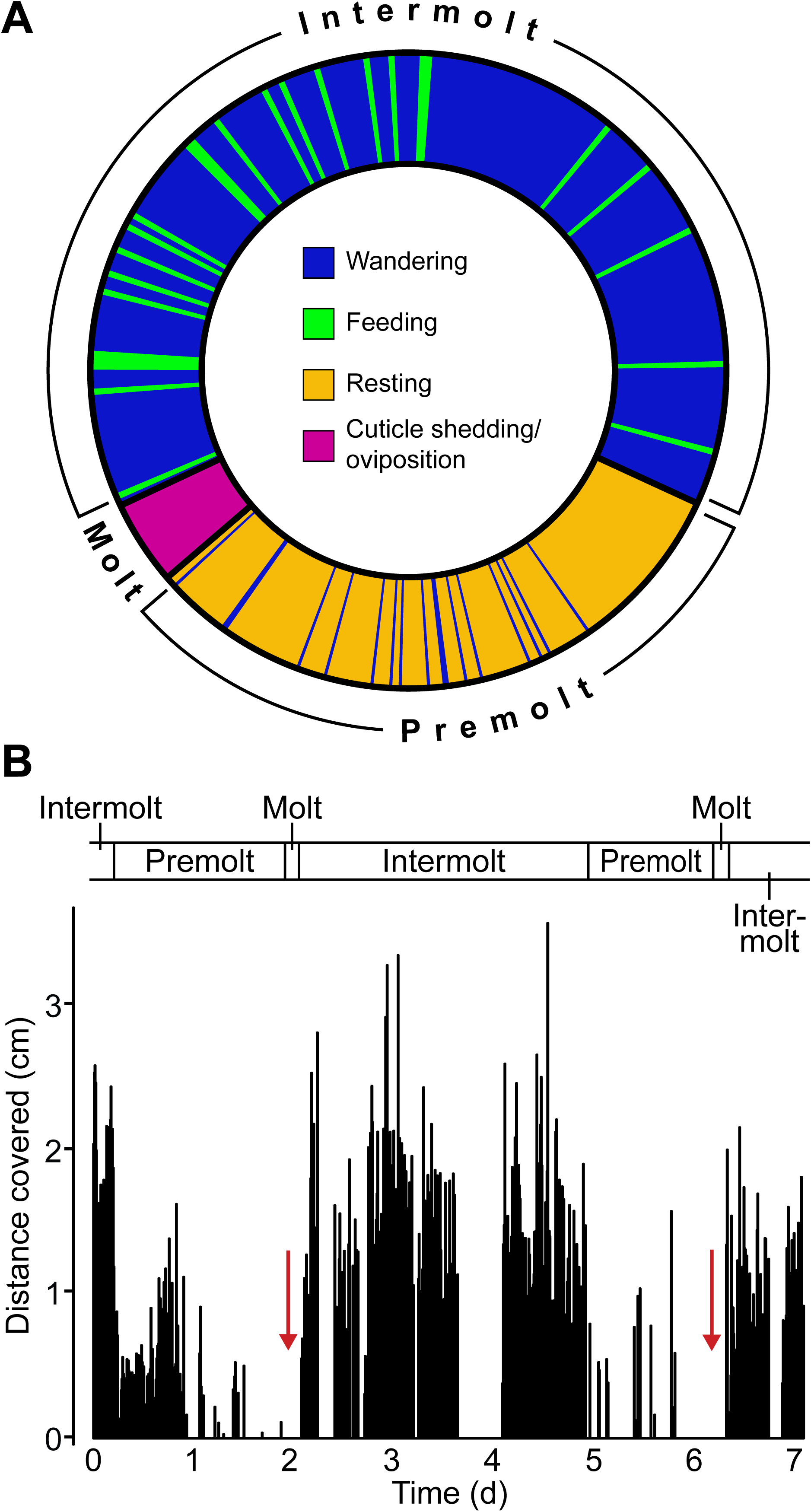
The ecdysial cycle is structured by recurring behavioral sequences. (A) Behavior combinations follow a consistent, repeating pattern across ecdysial cycles. Proportions represent the average across 10 individuals. The duration of feeding and wandering bouts is representative of typical values. No differences in behavior state combinations or stage durations were observed across lighting conditions. (B) Representative 7-day locomotor activity diagram from a single animal, starting 2 days after molt completion. Each peak corresponds to 10 min of activity. Red arrows indicate the onset of molt stages. The analysis focused on the interval between the two red arrows (one complete ecdysial cycle). The bar on top indicates the ecdysial stages: intermolt (INT), premolt (PRM), and molt (MLT).

**Table 1.** Behavioral states and their occurrence across ecdysial stages in *Hypsibius exemplaris*. Symbols: +, present; −, absent.

| Behavior | Description | Intermolt (INT) | Premolt (PRM) | Molt (MLT) |
| --- | --- | --- | --- | --- |
| Wandering (Movie S1) | Active locomotion across the substrate | + | + | – |
| Feeding (Movie S2) | Feeding on algae, either while slowly walking or crawling on algal clusters | + | – | – |
| Resting (Movie S3) | Complete behavioral quiescence, or subtle body and head twitches without locomotion | – | + | – |
| Cuticle shedding and oviposition (Movie S4) | Shedding of the old cuticle during ecdysis, with eggs deposited within the shed cuticle | – | – | + |

The macro-behavioral architecture of the tardigrade is organized into three distinct ecdysial stages: intermolt, premolt, and molt (Fig. 2B). The **intermolt (INT)** stage is characterized by alternating bouts of active wandering and feeding, lasting 2–4 days and comprising the majority of the total cycle (63.8%; *n* = 10). This stage exhibits the highest overall locomotor activity and represents the primary window for foraging and environmental exploration. The subsequent **premolt (PRM)** stage lasts 1–2 days (31.8% of the cycle), during which animals cease feeding entirely and exhibit prolonged resting bouts (ranging from 20 min to 8 h) punctuated by transient, wandering episodes (Fig. 2A). The **molt (MLT)** stage lasts 2– 4 h (4.4% of the cycle) and is exclusively defined by active cuticle shedding and oviposition (Movie S4). Crucially, the fundamental behavioral state combinations, overall cycle durations, and individual stage lengths do not differ across the three lighting regimes.

### Light–dark cycles modulate daily locomotor waveform shape

Daily median locomotor activity profiles reveal a distinct time-of-day organization under LD conditions. Analysis spanning the full ecdysial cycle demonstrates a prominent activity peak during the late dark phase (ZT19; Fig. 3A). Restricting the analysis exclusively to the highly active intermolt stage reveals a pronounced bimodal pattern, with peaks occurring at ZT8 and ZT19 (Fig. 3B). In contrast, individuals maintained in constant darkness (DD) exhibit a unimodal profile dominated by a single peak near projected ZT9 (Fig. 3C, D). This divergence suggests that the environmental LD cycle shapes the baseline locomotor waveform, potentially through direct photic masking and/or entrainment of endogenous oscillatory processes. A sensitivity analysis using pooled medians yields smoother behavioral profiles and confirms a minor peak at ZT8 in the overall LD profile (Fig. S1A), further supporting the robustness of the bimodal pattern (Fig. 3A, B), while intermolt LD and DD profiles remain structurally unchanged (Fig. S1B–D).

**Fig. 3.**
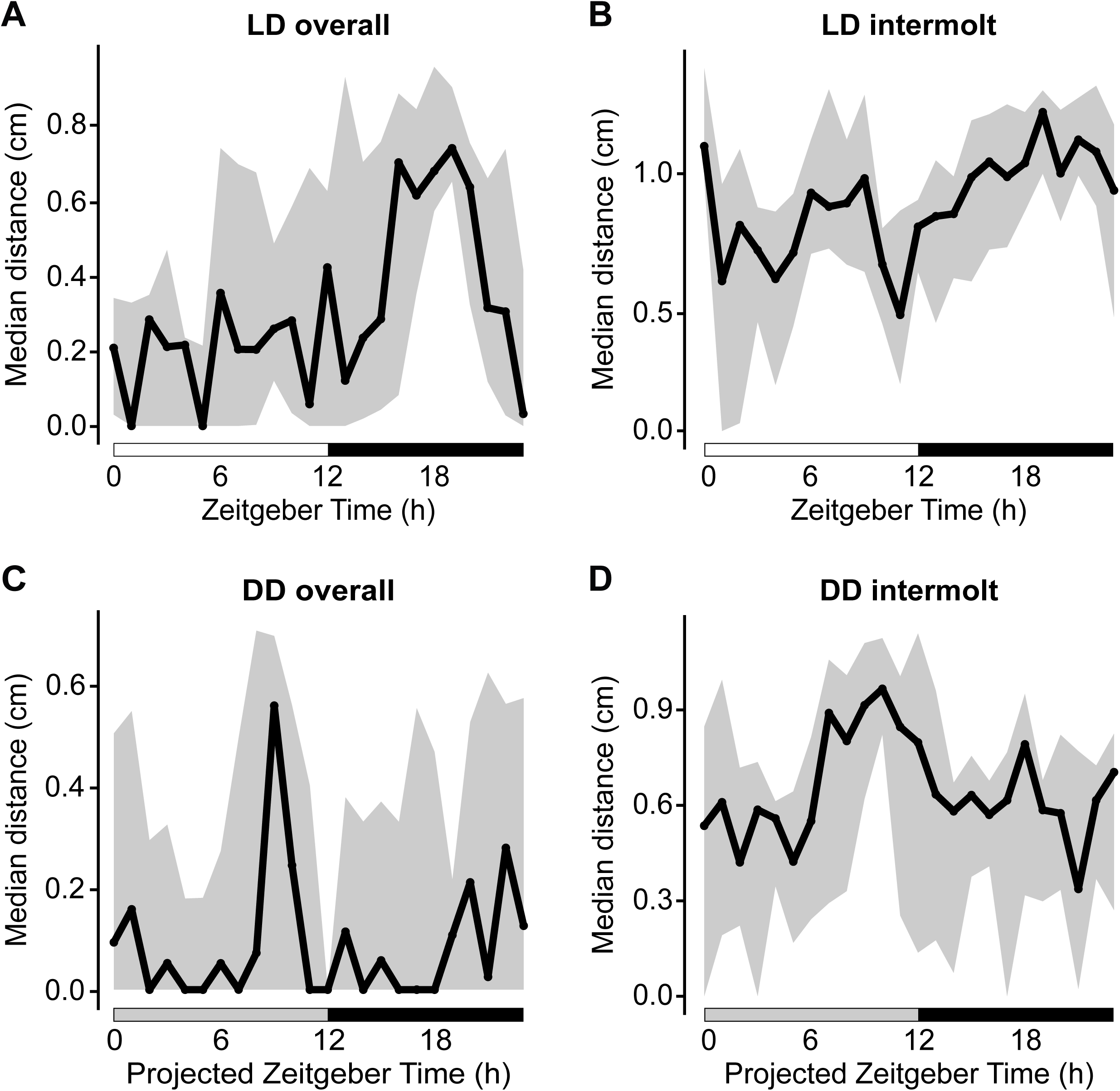
Daily activity profiles reveal time-of-day organization under LD and DD conditions. Median daily locomotor activity profiles are shown for (A) the full ecdysial cycle and (B) the intermolt (INT) stage under LD. A bimodal pattern is evident in the INT stage, with peaks near ZT8 and ZT19. (C) Under DD, the median profile shows a single peak near projected ZT9 over the full cycle. (D) Similarly, during the INT stage under DD, a single peak is observed near projected ZT9. Curves represent the median of per-animal median activity values at each hourly ZT/projZT bin (each animal weighted equally); shaded ribbons indicate the interquartile range across individuals. White and black horizontal bars indicate light and dark phases in LD panels; grey horizontal bars indicate the projected light–dark schedule in DD panels. Light/dark preference was assessed using paired Wilcoxon signed-rank tests (LD overall: one-sided Dark > Light, *p* = 0.06, *n* = 10; LD INT: *p* = 0.08, *n* = 10).

To statistically validate population-level time-of-day organization, we fitted cyclic generalized additive mixed models (GAMMs) with individual identity included as a random intercept. Both the LD profile (effective degrees of freedom [edf] = 5.27, *p* < 2 × 10⁻¹⁶) and the entrained DD profile (edf = 4.27, *p* = 0.001) exhibit significant time-of-day dependencies, confirming structured daily locomotor modulation even in the absence of external light. In contrast, naïve-DD animals lack a significant population-level dependence on a shared clock-time reference (edf ≈ 0, *p* = 0.76), consistent with the absence of synchronized population phase alignment prior to recording.

### Photic preference varies among individuals rather than at the population level

Despite clear time-of-day modulation of locomotor activity, population-level preference for either the light or dark phase is weak. A two-sided paired Wilcoxon signed-rank test reveals no significant difference in median intermolt activity between the light (ZT0–ZT11) and dark (ZT12–ZT23) phases (*p* = 0.16). However, a corresponding one-sided test indicates a trend toward greater dark-phase activity (Dark > Light, *p* = 0.08), with 7 of 10 individuals exhibiting higher cumulative activity during the dark phase (Fig. 4). This individual-level tendency is consistent with the dominant dark-phase peak observed in the population activity profile (Fig. 3A, B), suggesting that although a population-wide phase preference is not statistically significant, many individuals display a bias toward greater activity during the dark phase.

**Fig. 4.**
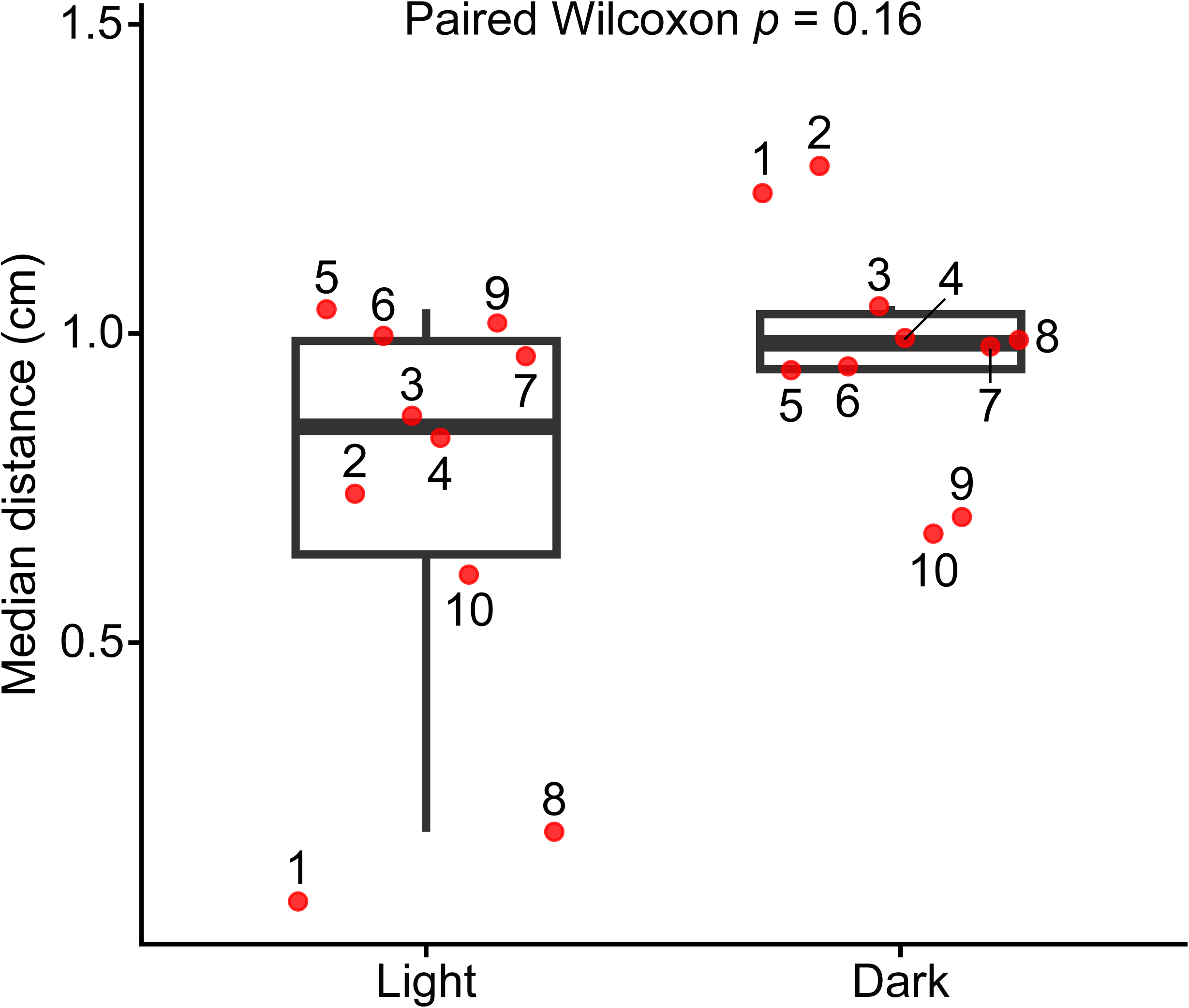
Individual variation in light/dark activity preference under LD. Each red dot represents the median intermolt locomotor activity in the dark (ZT12–ZT23) or light (ZT0– ZT11) phase for a single animal. Although 7 of 10 individuals exhibited higher activity in the dark, no significant population-level difference was detected (two-sided paired Wilcoxon signed-rank test, *p* = 0.16, *n* = 10). A one-sided test revealed a trend toward higher dark-phase activity (*p* = 0.08). For each individual, 10-min activity bins were normalized and assigned to light or dark phases. Horizontal black bars denote the median across individuals. Numbers adjacent to each dot indicate individual experiment IDs.

### Temporal partitioning of specific behaviors disrupted in constant darkness

We next examined the temporal distribution of individual behavioral states throughout the 24-h cycle. Under standard LD conditions, feeding and wandering exhibit distinct, complementary time-of-day organization: wandering peaks during late morning and mid-evening, whereas feeding occurs most frequently during the early light phase and around midday (Fig. 5A, B). Repeated-measures ANOVA (RM-ANOVA) and permutation tests confirm a significant time-of-day dependence for feeding behavior (*p* = 0.0003 and *p* = 0.0006, respectively), whereas wandering exhibits a weaker, non-significant trend (*p* = 0.08 for both tests). Resting and cuticle shedding/oviposition show no significant time-of-day dependence (*p* > 0.4 in both tests), although cuticle shedding is descriptively clustered around the light-on transition (ZT0) in the median profile (Fig. 5C, D). When quantified as overall behavioral proportions, wandering occurs 5% more frequently during the dark phase, whereas feeding and oviposition occur 20% more frequently during the light phase.

**Fig. 5.**
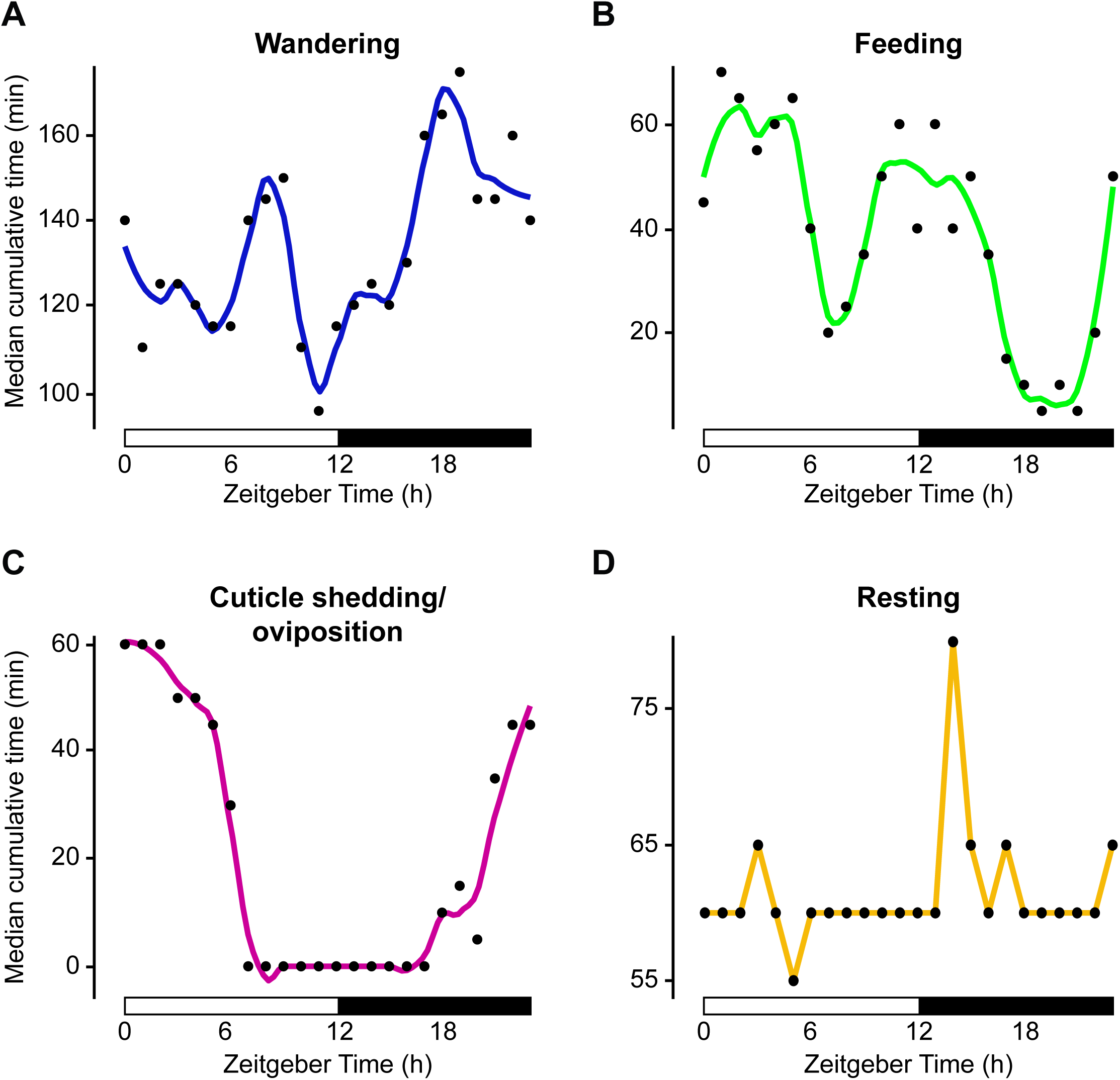
Time-of-day organization of behaviors under LD. The median number of cumulative minutes per hour spent in each behavioral category is plotted against ZT (0–23) across 10 LD animals. (A) Wandering shows two daily maxima near ZT9 and ZT19 (RM-ANOVA, *p* = 0.08). (B) Feeding peaks near ZT2 and ZT12 (RM-ANOVA, *p* < 0.001). (C) Cuticle shedding/oviposition is elevated near lights-on (ZT0) but shows no significant time-of-day dependence (RM-ANOVA *p* = 0.4, permutation test, *p* = 0.4). (D) Resting shows minimal time-of-day structure and is not significantly dependent on ZT (RM-ANOVA, *p* = 0.4, permutation test, *p* = 0.4). Curves in (A–C) were smoothed using LOESS (span = 0.3) for visualization. White and black horizontal bars indicate light and dark phases of the light–dark cycle.

In contrast, the temporal organization of both wandering and feeding is markedly degraded under constant darkness (Fig. S2A, B). Neither behavior retains significant time-of-day dependence in DD (RM-ANOVA and permutation tests, *p* > 0.1), suggesting that the precise temporal partitioning of these behavioral states depends on the presence of an environmental light–dark cycle and is not maintained by the endogenous oscillator alone.

### Locomotor rhythmicity persists in the absence of photic entrainment

To rigorously test for the presence of endogenous circadian rhythmicity, we applied a fixed-period 24-h cosinor model to detrended intermolt activity data. Individuals were classified as rhythmic based on a significant zero-amplitude F-test, with an *R²* > 0.08 threshold used as an additional criterion for detectable rhythmicity. Individual locomotor rhythmicity was detected across all three experimental cohorts: 6 of 10 individuals in LD, 7 of 11in DD, and 7 of 10 in naïve-DD met the rhythmicity criterion (Table S1).

The detection of clear daily rhythms in naïve-DD animals supports the presence of an endogenous circadian-like oscillator that regulates locomotor activity independently of prior light exposure. Although median rhythm amplitude (*A*) and *R²* values are numerically highest under LD conditions (Table S1), these differences are not statistically significant across the three environments (Kruskal–Wallis test: *A*, *p* = 0.18; *R²*, *p* = 0.31). These findings suggest that although environmental light may strengthen the expression of locomotor rhythms, it is not required for their generation.

### Phase heterogeneity accompanied by conserved circadian time-dependent modulation

Despite the presence of robust individual rhythms, absolute acrophases are broadly distributed around the circular phase space in both LD and DD conditions, showing no population-level phase synchronization (Rayleigh test, *p* > 0.05; Fig. S3). To isolate the underlying structure of daily modulation from this phase heterogeneity, we aligned individual activity profiles to each animal’s respective circadian time (CT), defining the individual cosinor acrophase as CT0.

The resulting CT-aligned median waveforms reveal a conserved bowl-shaped daily modulation pattern across all environmental cohorts, characterized by a reduction in activity around mid-CT and an increase near the aligned acrophase (Fig. 6). Pairwise comparisons using Pearson correlation and animal-level bootstrap resampling demonstrate significant structural similarity among the waveforms:

- **LD vs. DD:** *r =* 0.64, 95% CI [0.46, 0.71]
- **LD vs. naïve-DD:** *r =* 0.57, 95% CI [0.35, 0.68]
- **DD vs. naïve-DD:** *r =* 0.44, 95% CI [0.21, 0.60] (Table 2).

**Fig. 6.**
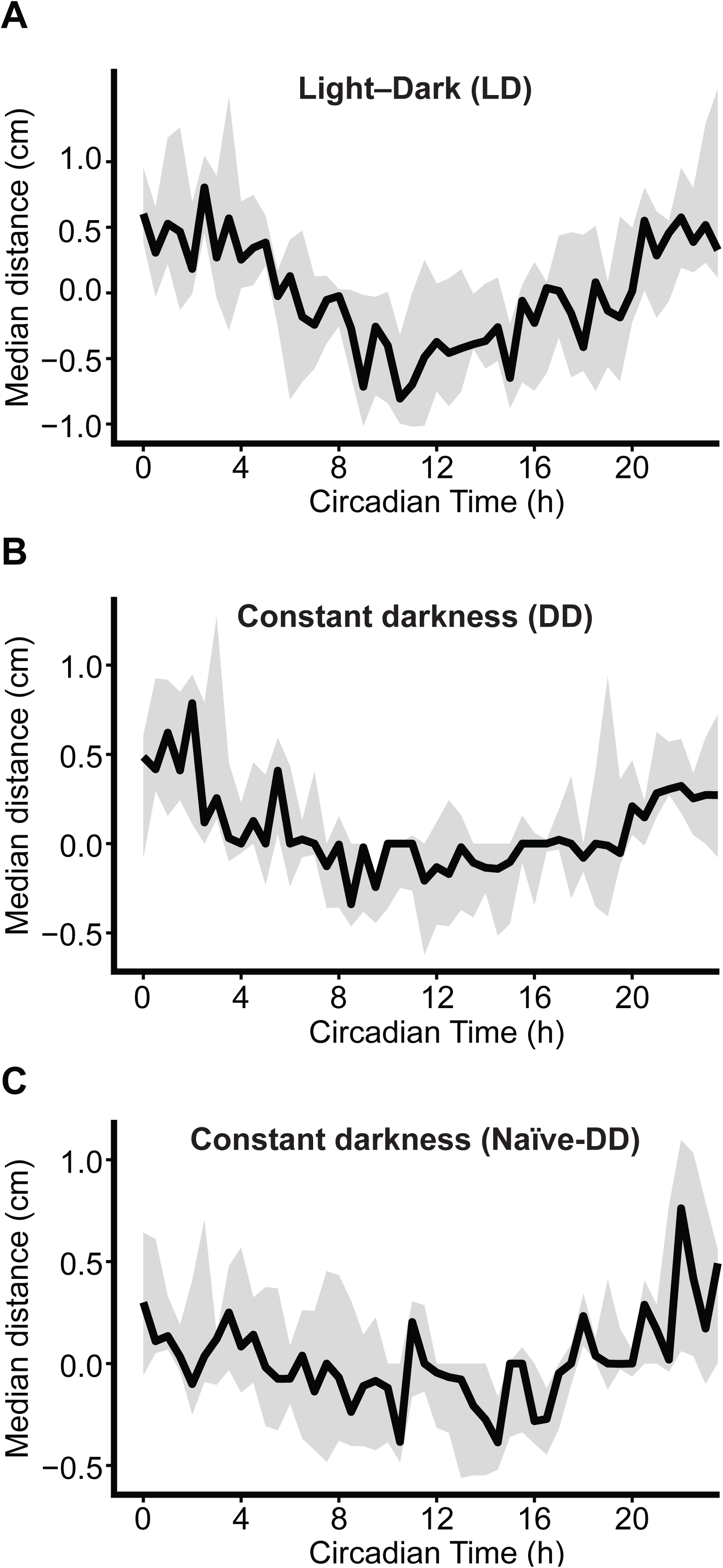
Circadian time-aligned median activity waveforms reveal conserved circadian-like modulation across lighting conditions. CT-aligned median waveforms are shown for (A) LD (*n* = 6), (B) DD (*n* = 7), and (C) naïve-DD (*n* = 7). Individuals were classified as rhythmic using the hourly-binned detrended 24-h cosinor fitting criterion (*R^2^* > 0.08; see Methods). Waveforms were constructed from detrended 10-min activity residuals, mapped to circadian time (CT) using each individual’s 24-h cosinor acrophase (CT0), and summarized in 30-min CT bins. Solid black lines represent the median across individuals; shaded ribbons indicate 95% animal-level bootstrap confidence intervals. Waveform similarity and amplitude differences are quantified in Table 2. The consistent shape of the waveforms across conditions, despite differences in amplitude, indicates a conserved circadian-time-dependent activity pattern.

**Table 2.** Circadian time-aligned waveform shape and amplitude comparisons across lighting conditions. Waveforms were constructed from detrended 10-min activity residuals aligned to circadian time (CT) using individual 24-h cosinor acrophases. Pearson correlation assessed waveform shape similarity, whereas amplitude differences quantified the strength of daily modulation. Comparisons are based on animal-level bootstrap resampling (*B* = 2,000 iterations). The 95% CIs include zero for DD vs. naïve-DD, indicating no detectable difference in amplitude between these conditions.

| Comparison | Shape similarity (95% bootstrap CI) | Amplitude difference (95% bootstrap CI) |
| --- | --- | --- |
| LD vs DD | [0.46, 0.71] | [0.12, 1.02] |
| LD vs naïve-DD | [0.35, 0.68] | [0.12, 1.03] |
| DD vs naïve-DD | [0.21, 0.60] | [–0.51, 0.47] |

Parallel Spearman rank correlations confirm these patterns (e.g., LD vs. DD: *ρ* = 0.77, 95% CI [0.51, 0.76]), validating the mathematical robustness of the shared waveform shape.

The baseline modulation amplitude—quantified as the peak-to-trough distance of the median waveform—is significantly greater in LD (1.61, 95% CI [1.39, 1.99]) than in entrained DD (1.12, 95% CI [0.75, 1.47]) and naïve-DD (1.14, 95% CI [0.71, 1.47]). Bootstrap-estimated amplitude differences between LD and DD (0.48, 95% CI [0.12, 1.02]) and between LD and naïve-DD (0.46, 95% CI [0.12, 1.03]) are consistently positive, whereas the two constant darkness conditions do not differ from each other (0.02, 95% CI [−0.51, 0.47]) (Table 2).

Furthermore, total cumulative intermolt activity is significantly elevated in LD relative to both DD regimes (Benjamini-Hochberg-adjusted pairwise Wilcoxon test, *p* = 0.014 for both), while the two DD conditions were statistically indistinguishable (*p* > 0.1). Taken together, these metrics indicate that environmental light cycles enhance both the amplitude of circadian modulation and overall baseline activity levels but do not generate the underlying daily modulation itself.

To evaluate the persistence of this endogenous modulation, we extended the analysis to 48-h CT-aligned waveforms (Fig. S4). A fixed-period 24-h cosinor model fitted to the group median waveform confirms that rhythm strength and amplitude are highest in LD (*A* = 0.63, *R²* = 0.67), followed by DD (*A* = 0.26, *R²* = 0.51) and naïve-DD (*A* = 0.20, *R²* = 0.30), validating sustained rhythmicity across multiple subjective days. Finally, a descriptive analysis of the raw, non-aligned 48-h population profile in LD independently supports the presence of a sustained 24-h rhythm component (cosinor *A* = 0.13, *R²* = 0.27; Fig. S5).

## DISCUSSION

This study provides evidence consistent with an endogenous, circadian-like timing mechanism in tardigrades and characterizes how daily locomotor activity is integrated with the ecdysial cycle of *Hypsibius exemplaris*. By tracking behavior continuously across multiple days, we revealed a highly reproducible, three-stage ecdysial sequence—intermolt (INT), premolt (PRM), and molt (MLT)—that dictates overall behavioral states independently across lighting conditions. This staging framework may improve the design and interpretation of future work, including gene-expression studies, pharmacological assays, and stress-tolerance experiments, by encouraging researchers to account for behavioral and physiological changes across the ecdysial cycle.

### Molt-dependent gating: Ecdysial regulation of activity and quiescence

A major finding is that *H. exemplaris* rest–activity patterns are governed by ecdysial progression rather than a rigid daily rest phase. During the highly active intermolt stage, individuals exhibit sustained, continuous foraging and exploration across multiple days. The transition to the premolt stage is marked by a dramatic behavioral shift: animals cease feeding and display prolonged resting bouts.

This PRM quiescence consistently emerges approximately 24 h prior to cuticle shedding, bearing a striking resemblance to the sleep-like “lethargus” state observed during *Caenorhabditis elegans* molts (Trojanowski and Raizen, 2016; Raizen et al., 2008; Iwanir et al., 2013). In *C. elegans*, molt-dependent/developmental transitions and quiescent states are tightly regulated by nuclear receptor pathways, such as *nhr-23* (Hiroki and Yoshitane, 2024). Given that ecdysteroid levels fluctuate dynamically across the tardigrade ecdysial cycle (Yamakawa and Hejnol, 2024), endocrine signaling is a highly plausible driver of this premolt resting state.

Although resolution constraints prevented direct observation of the morphological simplex stage, defined by stylet discharge and mouth closure (Guidetti, 2025), the cessation of feeding during PRM and MLT is consistent with a transition toward this stage. Because resting likely precedes stylet discharge, simplex may begin after, rather than coincide with, PRM onset. Resting therefore appears to be associated primarily with ecdysial stage, whereas active behaviors during INT remain subject to temporal regulation.

### The circadian “envelope” and photic waveform shaping

When isolating the intermolt stage, our data reveal an endogenous circadian-like signal operating beneath a bout-dominated behavioral repertoire. Rather than enforcing a rigid, clock-timed daily schedule, this clock functions primarily as an activity envelope, modulating the probability and intensity of locomotor bouts over a ∼24-h cycle. This type of behavioral organization may be an evolutionarily widespread strategy among metazoans whose immediate behaviors are highly stochastic and state-dependent (Helfrich-Förster, 2000; Schwartz and Klerman, 2019).

Under light–dark cycles (LD), the population-level activity profile is distinctly bimodal, characterized by a minor morning peak and a dominant late-dark phase peak. This bimodal distribution disappears in constant darkness (DD), shifting to a unimodal profile centered around the projected daytime. This phenomenon mirrors behavioral dynamics observed in *Drosophila melanogaster* and sea anemones, where complex waveforms arise from the interplay between an internal oscillator and direct light-dependent effects, or masking (Helfrich-Förster, 2000; Hendricks et al., 2012).

While light significantly enhances the amplitude of these rhythms and boosts overall intermolt activity levels, the persistence of robust 24-h rhythmicity in both entrained DD and naïve-DD individuals indicates that photic input shapes and amplifies the activity waveform but is not required for its generation.

### Phase heterogeneity and microhabitat adaptation

Intriguingly, despite using a parthenogenetic, genetically clonal line, we observed substantial inter-individual variability and a lack of population-level phase locking across all conditions. Absolute phases relative to zeitgeber time were highly heterogeneous. This structural divergence—where individuals maintain robust independent rhythms but do not align to a common population phase—suggests a flexible timing system.

For a microscopic organism inhabiting highly variable, fragmented microhabitats, tight population-level synchronization may be maladaptive. Instead, an uncoupled, individual-level phase strategy may facilitate opportunistic foraging or cryptic predator avoidance tailored to immediate microenvironmental cues (Moore et al., 1989; Xu et al., 2008; Ingram et al., 2016).

### Methodological insights and future directions

Quantifying behavior over extended timelines in a microscopic organism presents distinct technical trade-offs. Deep learning-based tracking via DeepLabCut proved highly robust at isolating individual trajectories amidst visual clutter from food sources, echoing recent successes in tardigrade gait classification (Ramahefarivo et al., 2026). However, our long-term recordings were fundamentally constrained to a single ecdysial cycle; maintaining sufficient algae was mandatory to stimulate feeding and trigger molting, yet excessive algal growth compromised tracking accuracy. Because individual baseline activity patterns exhibited slow, ecdysial state-dependent drift, mathematical detrending was essential to isolate the ∼24-h circadian modulation from molt-dependent baseline shifts.

While these findings establish a foundational behavioral framework for tardigrade chronobiology, several canonical properties of a formal circadian system remain to be verified (Vitaterna et al., 2001). Due to sample size constraints and individual phase heterogeneity, future studies employing larger cohorts, phase-shifting assays, and temperature-compensation experiments will be critical to confirm a canonical circadian clock. Furthermore, identifying the spatial expression of candidate clock coupling factors, such as the recently mapped pigment-dispersing factor genes (*He-pdf-1*, *-2*, and *-3*; Mayer et al., 2015; Dutta et al., 2025), will provide a vital molecular bridge to understanding the evolution of biological timing within the Ecdysozoa.

## Conclusions

In conclusion, this study provides evidence that *Hypsibius exemplaris* possesses an endogenous, circadian-like timing mechanism that modulates locomotor activity within a strictly regulated ecdysial framework. Rather than driving a rigid behavioral schedule, this internal clock operates as a decentralized activity “envelope” that modulates the probability and intensity of active states during the intermolt stage. While environmental light–dark cycles actively shape and amplify the behavioral waveform, photic input is not mandatory for its expression. The striking coexistence of robust individual rhythmicity and population-level phase heterogeneity points to a highly flexible, opportunistic timing strategy uniquely suited to cryptic, variable microhabitats. By mapping these dynamics, this work establishes tardigrades as a powerful and tractable model for ecdysozoan chronobiology, opening a new window into how daily timing systems intersect with ecdysial cycles and evolve across diverse life histories.

## Acknowledgements

We thank the members of the Research Training Group “Biological Clocks on Multiple Time Scales” (GRK 2749/1) for constructive discussions. We particularly thank Sandra Treffkorn for discussions of ecdysial stages; Huleg Zolmon and Anna Schneider for critical feedback on data analysis and manuscript preparation; and Jenny Plath for technical assistance with imaging and guidance on using the DeepLabCut tracking software.

## Author contributions

Conceptualization: G.M., L.H., B.C.U., N.F.; Data acquisition and behavioral analysis: B.C.U., L.H.; Mathematical analysis: N.F., E.F., B.C.U.; Visualization: B.C.U., G.M.; Funding acquisition: G.M., E.F.; Writing – original draft: B.C.U., G.M.; Writing – review & editing: B.C.U., N.F., L.H., E.F., G.M.

## Data availability

Raw tracking data and custom scripts are deposited in Zenodo under accession number [DOI].

## SUPPLEMENTARY MATERIAL

**Table S1.** Cosinor24 fitting results for individual experiments. Rhythmicity was defined as *R^2^* > 0.08, and individuals (#) meeting this criterion are marked with an asterisk (*). Rhythmic individuals were included in circadian time (CT)-aligned waveform construction and bootstrap analyses. Amplitude is reported in µm·s⁻¹ (mean speed). Zero-amplitude *p* values indicate the statistical significance of deviation from a zero-amplitude (flat) cosinor model.

| # | Condition | Amplitude ($\mu\text{m}\cdot\text{s}^{-1}$) | $R^2$ | Zero-amplitude $p$ |
| --- | --- | --- | --- | --- |
| 1 | LD | 3.46 | 0.316* | < 0.001 |
| 2 | LD | 2.10 | 0.140* | 0.009 |
| 3 | LD | 1.13 | 0.060 | 0.186 |
| 4 | LD | 1.04 | 0.107* | 0.036 |
| 5 | LD | 0.21 | 0.008 | 0.809 |
| 6 | LD | 3.15 | 0.312* | < 0.001 |
| 7 | LD | 0.83 | 0.031 | 0.516 |
| 8 | LD | 2.27 | 0.290* | < 0.001 |
| 9 | LD | 0.20 | 0.003 | 0.912 |
| 10 | LD | 4.34 | 0.360* | < 0.001 |
| 1 | DD | 2.78 | 0.380* | < 0.001 |
| 2 | DD | 1.51 | 0.230* | < 0.001 |
| 3 | DD | 1.11 | 0.170* | 0.006 |
| 4 | DD | 1.08 | 0.083* | 0.090 |
| 5 | DD | 0.37 | 0.020 | 0.505 |
| 6 | DD | 1.02 | 0.136* | 0.014 |
| 7 | DD | 1.13 | 0.031 | 0.390 |
| 8 | DD | 3.69 | 0.550* | < 0.001 |
| 9 | DD | 0.65 | 0.037 | 0.346 |
| 10 | DD | 1.75 | 0.240* | 0.001 |
| 11 | DD | 0.76 | 0.045 | 0.258 |
| 1 | naïve-DD | 0.50 | 0.042 | 0.290 |
| 2 | naïve-DD | 0.75 | 0.103* | 0.055 |
| 3 | naïve-DD | 1.27 | 0.108* | 0.067 |
| 4 | naïve-DD | 1.26 | 0.216* | < 0.001 |
| 5 | naïve-DD | 1.41 | 0.114* | 0.048 |
| 6 | naïve-DD | 2.28 | 0.223* | < 0.001 |
| 7 | naïve-DD | 0.76 | 0.061 | 0.200 |
| 8 | naïve-DD | 0.48 | 0.022 | 0.530 |
| 9 | naïve-DD | 1.98 | 0.261* | 0.001 |
| 10 | naïve-DD | 1.44 | 0.179* | 0.003 |

**Fig. S1. Daily activity profiles under LD and DD with unweighted averaging.** Median daily locomotor activity profiles are shown for (A, C) the full ecdysial cycle and (B, D) the intermolt (INT) stage under LD and DD. All 10-min activity bins were included in the median calculation, with individuals weighted by the number of observations per ZT bin (i.e., without equal weighting of individuals). In LD, a prominent peak occurred at ZT8–ZT9 and another at ZT19 in the full cycle and INT stage. In DD, a single dominant peak was observed near the projected ZT9 for both the full cycle and the INT stage. For each animal, activity bins were normalized. White, black, and grey horizontal bars indicate light, dark, and projected light phases of the light–dark cycle, respectively. Vertical lines represent the interquartile range across individuals. These profiles support the bimodality of the daily activity pattern under LD and the persistence of a single peak in DD, when individual contributions are not equally weighted.

**Fig. S2. Time-of-day organization of behaviors under DD.** The median number of cumulative minutes per hour spent in each behavioral category is plotted against projected ZT (0–23) across 11 DD animals. (A) Wandering exhibited a single peak near projected ZT13. (B) Feeding showed peaks near projected ZTs: 7, 11, and 20. (C) Cuticle shedding/oviposition peaked at projected ZT5. (D) Resting showed minimal variation across time. None of the behaviors exhibited significant time-of-day dependence (RM-ANOVA and permutation tests, *p* > 0.1). For visualization, curves in (A–C) were smoothed using LOESS (span = 0.3). Grey and black horizontal bars indicate projected light and dark phases of the light–dark cycle, respectively. The lack of significant time-of-day structure under DD supports the conclusion that temporal organization of these behaviors is reduced in constant darkness.

**Fig. S3. Acrophase distributions reveal no population-level phase synchronization.** Circular plots show the distribution of individual 24-h cosinor acrophases (black dots) expressed in ZT (LD), projected ZT (DD). (A) LD (*n* = 6, Rayleigh test *p* = 0.4), (B) DD (*n* = 7, *p* = 0.7). In all conditions, acrophases were broadly distributed with no significant clustering (Rayleigh test, *p* > 0.1), indicating the absence of population-level phase locking to the light– dark cycle. Numbers around the circle represent ZT for LD and projected ZT for DD conditions. This phase heterogeneity underscores the importance of using circadian time (CT) rather than absolute time (ZT) for waveform comparisons.

**Fig. S4. Circadian time-aligned median activity waveforms over two consecutive cycles.** CT-aligned median waveforms are shown over 48 h for rhythmic individuals in (A) LD (*n* = 6), (B) DD (*n* = 7), and (C) naïve-DD (*n* = 7). Detrended 10-min activity residuals were mapped to circadian time (CT) using each individual’s 24-h cosinor acrophase (CT0) and summarized in 30-min CT bins. Solid lines indicate the median across individuals; shaded ribbons show the interquartile range across individuals. Dashed lines mark the 24-h boundary. For descriptive comparison of waveform sinusoidality, a fixed-period 24-h cosinor fit to the 48-h group-median curve yielded: LD (amplitude *A* = 0.63, *R²* = 0.67), DD (*A* = 0.26, *R²* = 0.51), and naïve-DD (*A* = 0.20, *R²* = 0.30). The higher amplitude (*A*) and rhythm strength (*R²*) in LD indicate stronger rhythmicity, while the persistence of modulation across cycles supports the stability of the circadian component.

**Fig. S5. Descriptive analysis of LD intermolt locomotor activity over 48 hours.** Median intermolt locomotor activity is shown over 48 h for 10 LD animals. Activity was detrended using a 30-h rolling median, and the median across individuals was plotted. The signal exhibits a clear 24-h rhythm, confirmed by a fixed-period 24-h cosinor fit (*A* = 0.13, *R^2^* = 0.27). The grey ribbon indicates the interquartile range across individuals. The dashed line marks the 24-h boundary. This analysis provides a descriptive confirmation of circadian modulation in the LD condition, consistent with the findings from CT-aligned waveform analysis.

## MOVIE CAPTIONS

**Movie S1. Example locomotor activity tracking of *Hypsibius exemplaris*.** Locomotor behavior was tracked using DeepLabCut based on the central landmark of the animal. The video was recorded at 10 frames per second and covers 32 minutes. Tracking artifacts resulting in abrupt, spatially distant jumps were corrected by coordinate interpolation before subsequent analyses. The timestamp indicates real-world elapsed time.

**Movie S2. Example of feeding behavior while crawling on an algae cluster.** The video was recorded at 10 frames per second and magnified by a factor of 1.9. The corresponding 10-min locomotor activity bins for this behavior were assigned an activity value of 0. The timestamp indicates real-world elapsed time.

**Movie S3. Example of resting behavior.** The video was recorded at 10 frames per second and magnified by a factor of 1.9. The center point of the animal was tracked by DeepLabCut. The corresponding 10-min locomotor activity bins for this behavior were assigned an activity value of 0 when tracking was inaccurate due to the animal being covered by algae or curled up. The timestamp indicates real-world elapsed time.

**Movie S4. Example of cuticle shedding/oviposition behavior.** The video was recorded at 10 frames per second and magnified by a factor of 1.9. The center point of the animal was tracked by DeepLabCut. The corresponding 10-min locomotor activity bins for this behavior were assigned an activity value of 0. The timestamp indicates real-world elapsed time.

