## Supplementary Figure 1 for "Daily locomotor rhythms and ecdysial stage-dependent behaviors in the tardigrade *Hypsibius exemplaris*"

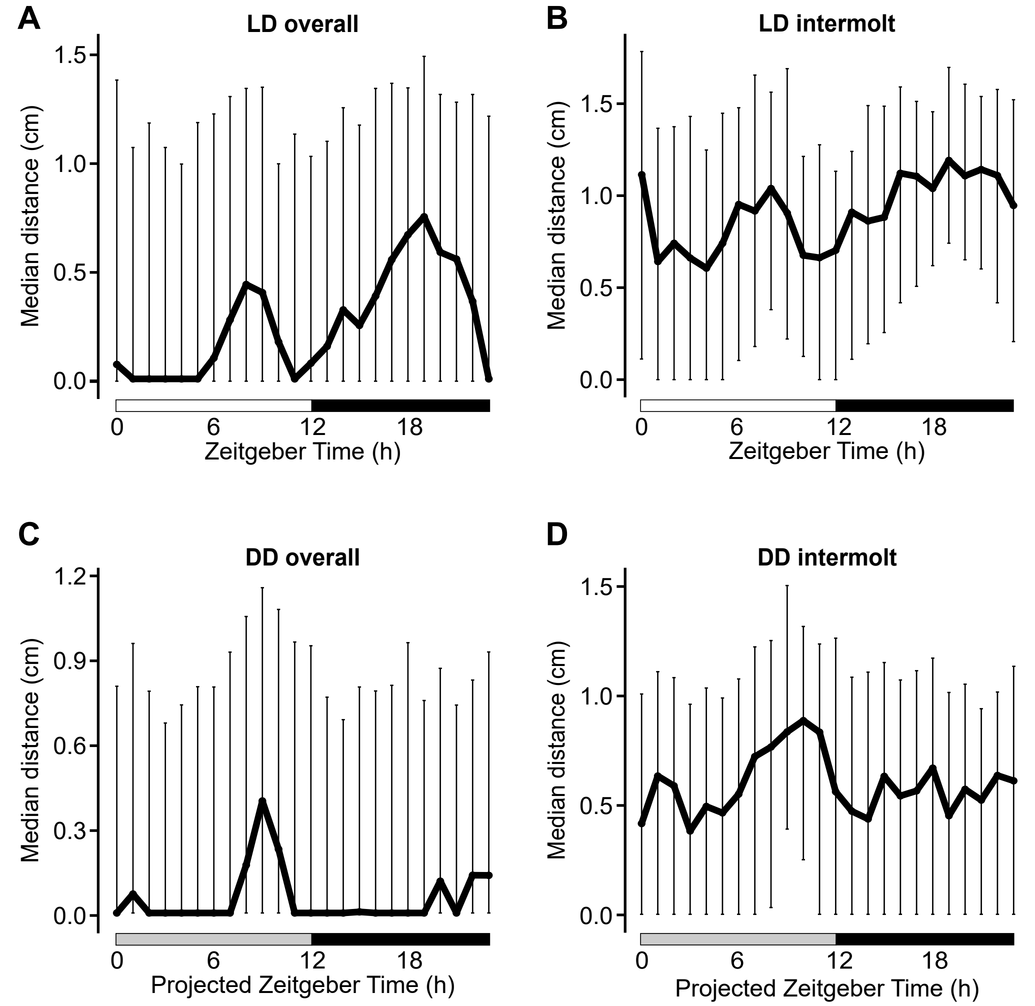


**Fig. S1. Daily activity profiles under LD and DD with unweighted averaging.** Median daily locomotor activity profiles are shown for (A, C) the full ecdysial cycle and (B, D) the intermolt (INT) stage under LD and DD. All 10-min activity bins were included in the median calculation, with individuals weighted by the number of observations per ZT bin (i.e., without equal weighting of individuals). In LD, a prominent peak occurred at ZT8–ZT9 and another at ZT19 in the full cycle and INT stage. In DD, a single dominant peak was observed near the projected ZT9 for both the full cycle and the INT stage. For each animal, activity bins were normalized. White, black, and grey horizontal bars indicate light, dark, and projected light phases of the light–dark cycle, respectively. Vertical lines represent the interquartile range across individuals. These profiles support the bimodality of the daily activity pattern under LD and the persistence of a single peak in DD, when individual contributions are not equally weighted.
