## Supplementary Figure 2 for "Daily locomotor rhythms and ecdysial stage-dependent behaviors in the tardigrade *Hypsibius exemplaris*"

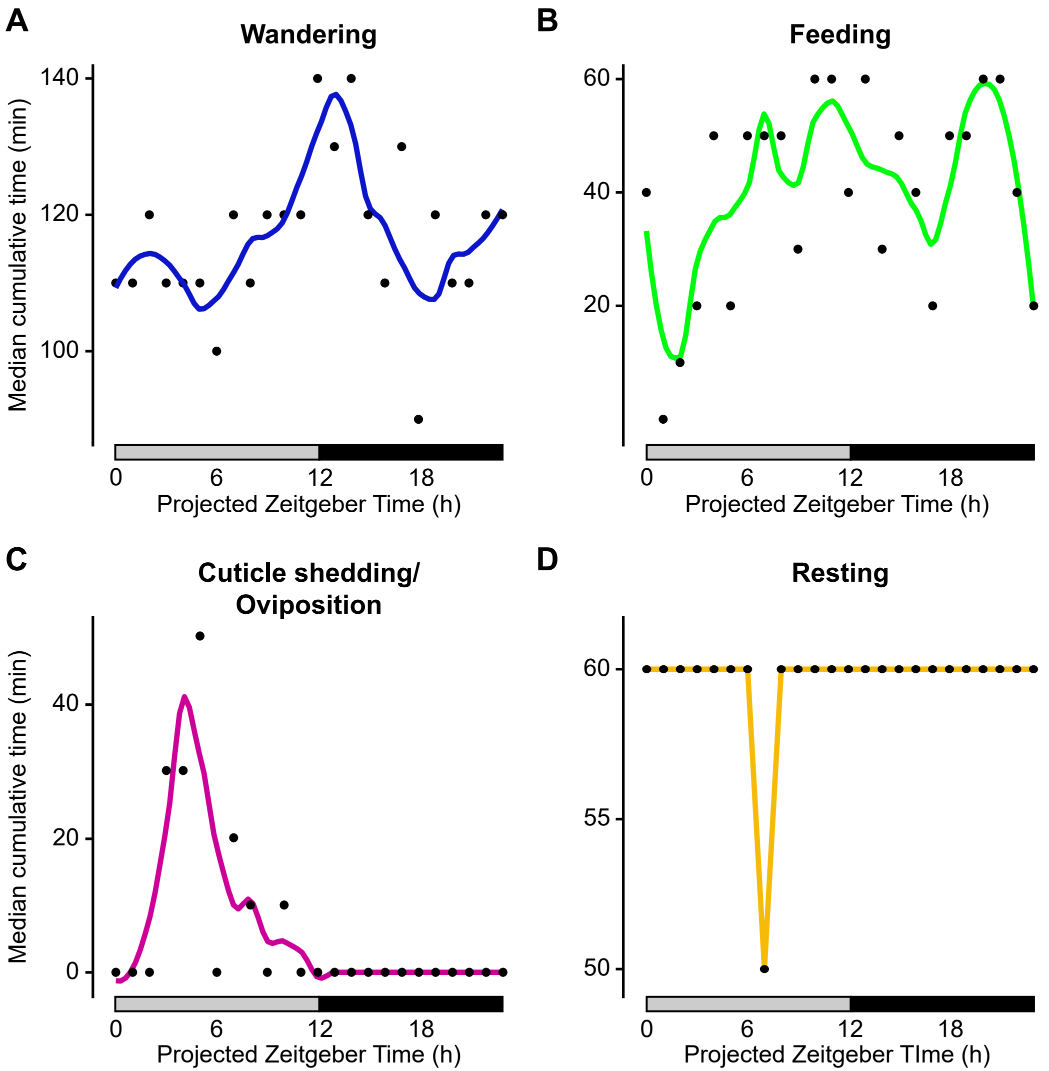


**Fig. S2. Time-of-day organization of behaviors under DD.** The median number of cumulative minutes per hour spent in each behavioral category is plotted against projected ZT (0–23) across 11 DD animals. (A) Wandering exhibited a single peak near projected ZT13. (B) Feeding showed peaks near projected ZTs: 7, 11, and 20. (C) Cuticle shedding/oviposition peaked at projected ZT5. (D) Resting showed minimal variation across time. None of the behaviors exhibited significant time-of-day dependence (RM-ANOVA and permutation tests, $p > 0.1$). For visualization, curves in (A–C) were smoothed using LOESS (span = 0.3). Grey and black horizontal bars indicate projected light and dark phases of the light–dark cycle, respectively. The lack of significant time-of-day structure under DD supports the conclusion that temporal organization of these behaviors is disrupted in constant darkness.
