## Supplementary Figure 3 for "Daily locomotor rhythms and ecdysial stage-dependent behaviors in the tardigrade *Hypsibius exemplaris*"

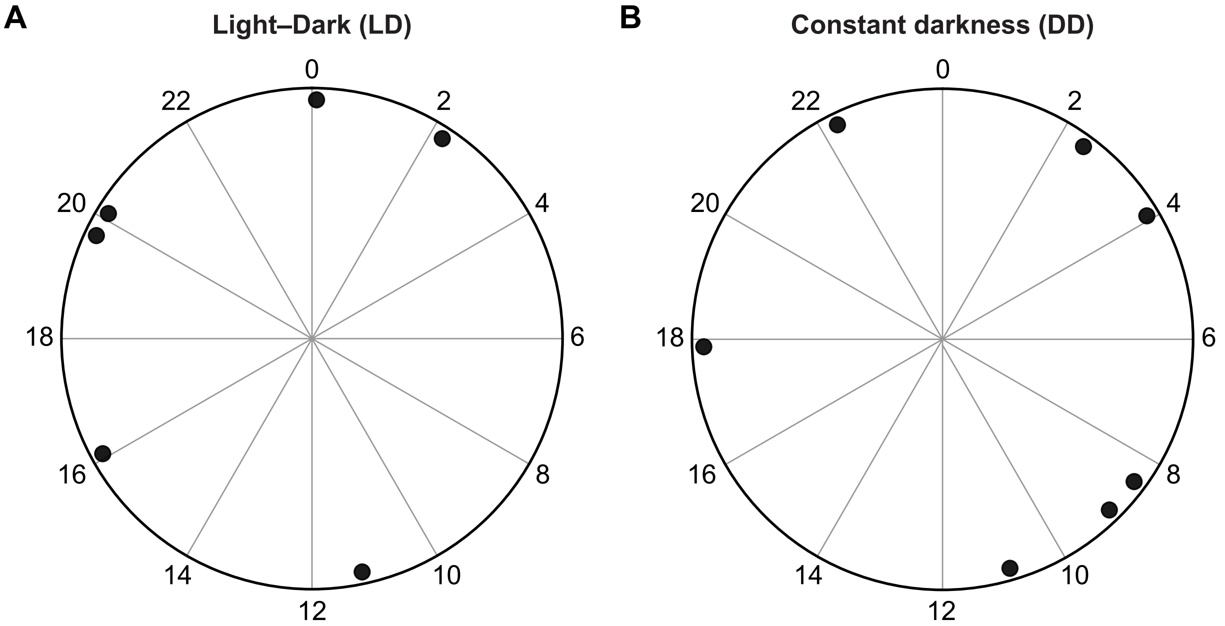


**Fig. S3. Acrophase distributions reveal no population-level phase synchronization.** Circular plots show the distribution of individual 24-h cosinor acrophases (black dots) expressed in ZT (LD), projected ZT (DD). (A) LD (n = 6, Rayleigh test $p = 0.4$), (B) DD (n = 7, $p = 0.7$). In all conditions, acrophases were broadly distributed with no significant clustering (Rayleigh test, $p > 0.1$), indicating the absence of population-level phase locking to the light–dark cycle. Numbers around the circle represent ZT for LD and projected ZT for DD conditions. This phase heterogeneity underscores the importance of using circadian time (CT) rather than absolute time (ZT) for waveform comparisons.
