## Supplementary Figure 4 for "Daily locomotor rhythms and ecdysial stage-dependent behaviors in the tardigrade *Hypsibius exemplaris*"

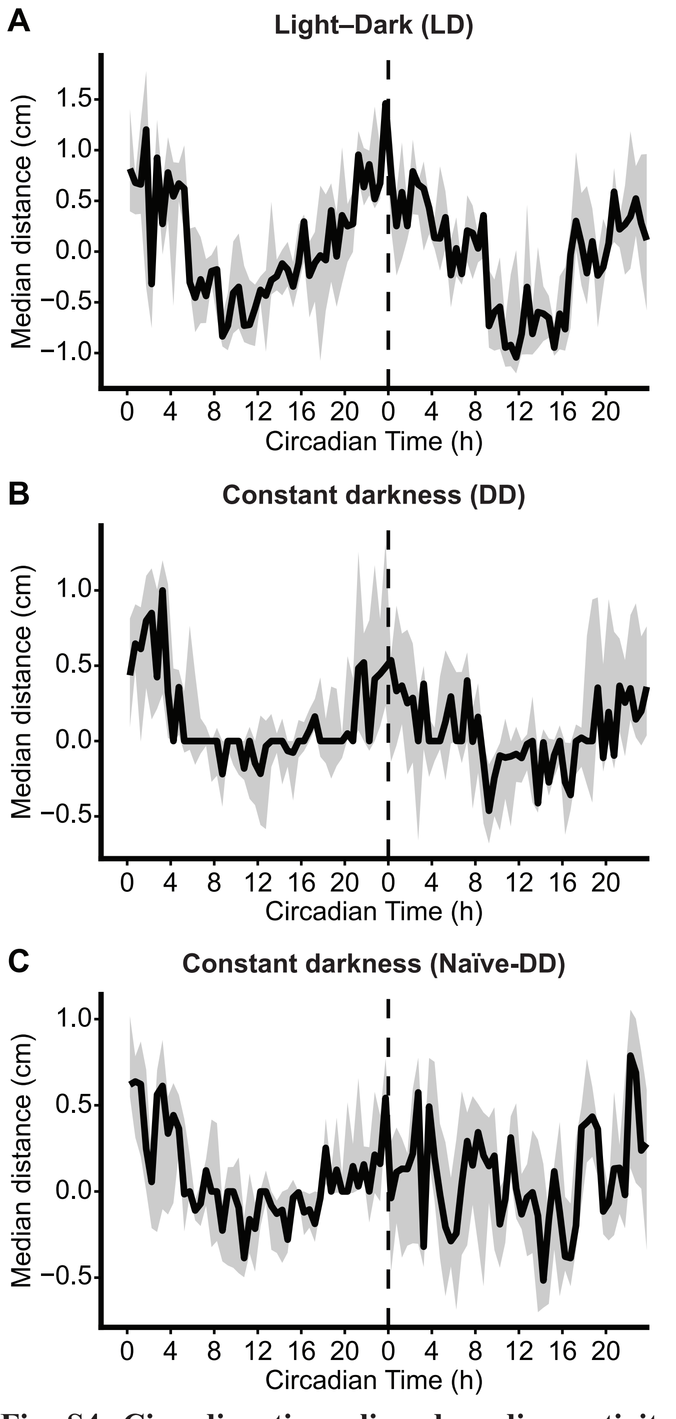


**Fig. S4. Circadian time-aligned median activity waveforms over two consecutive cycles.** CT-aligned median waveforms are shown over 48 h for rhythmic individuals in (A) LD ($n = 6$), (B) DD ($n = 7$), and (C) naïve-DD ($n = 7$). Detrended 10-min activity residuals were mapped to circadian time (CT) using each individual's 24-h cosinor acrophase (CT0) and summarized in 30-min CT bins. Solid lines indicate the median across individuals; shaded ribbons show the interquartile range across individuals. Dashed lines mark the 24-h boundary. For descriptive comparison of waveform sinusoidality, a fixed-period 24-h cosinor fit to the 48-h group-median curve yielded: LD (amplitude $A = 0.63$, $R^2 = 0.67$), DD ($A = 0.26$, $R^2 = 0.51$), and naïve-DD ($A = 0.20$, $R^2 = 0.30$). The higher amplitude ($A$) and rhythm strength ($R^2$) in LD indicate stronger rhythmicity, while the persistence of modulation across cycles supports the stability of the circadian component.
