## Supplementary Figure 5 for "Daily locomotor rhythms and ecdysial stage-dependent behaviors in the tardigrade *Hypsibius exemplaris*"

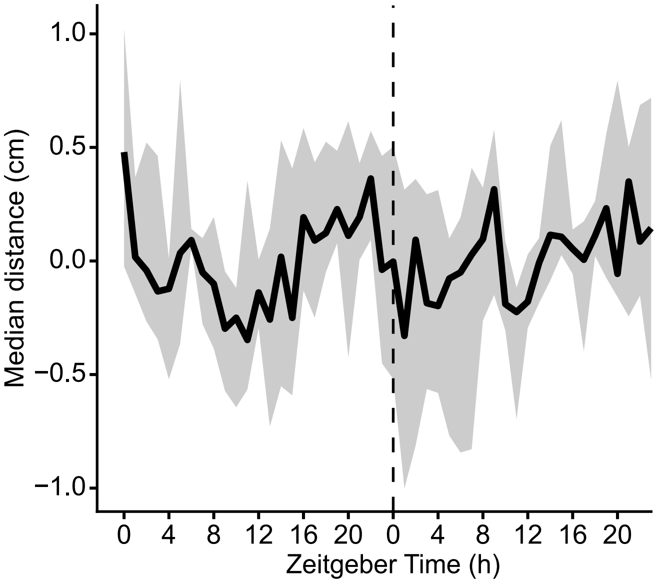


**Fig. S5. Descriptive analysis of LD intermolt locomotor activity over 48 hours.** Median intermolt locomotor activity is shown over 48 h for 10 LD animals. Activity was detrended using a 30-h rolling median, and the median across individuals was plotted. The signal exhibits a clear 24-h rhythm, confirmed by a fixed-period 24-h cosinor fit ($A$ = 0.13, $R2$ = 0.27). The grey ribbon indicates the interquartile range across individuals. The dashed line marks the 24-h boundary. This analysis provides a descriptive confirmation of circadian modulation in the LD condition, consistent with the findings from CT-aligned waveform analysis.
